# Biased evolution of pollen size after whole genome duplication

**DOI:** 10.64898/2026.11.13.766547

**Authors:** Joseph H. Williams

## Abstract

Cell size is an important functional attribute of single-celled organisms, and whole genome duplication (WGD) has long been hypothesized to double the volume of unicells. Pollen of flowering plants is effectively a single-celled organism, comprising the “vegetative cell” of the male gametophyte, which carries the male germline (two sperm cells or their precursor) within its cytoplasm. Instantaneous doubling of evolved pollen size is likely to be excessive, which should bias evolution in young polyploids to size reduction. Here, I test the hypotheses that the magnitude of the effect of WGD on pollen diameter should be doubled volume at inception, and less-than-doubled volume in evolved autotetraploids. Taking data from the literature, I compared 1*n* = 2*x* to 1*n* = 1*x* pollen diameters in 286 pairs of intraspecific autotetraploids and their diploid progenitors at four successive evolutionary ages: 1) diploid anthers producing some non-reduced pollen, 2) newly formed synthetic or spontaneous autotetraploids, 3) established autotetraploid cultivars, and 4) wild intraspecific autotetraploid cytotypes. Relative to 1*x* pollen, the volume of 2*x* non-reduced pollen was *at least doubled* at inception (31 species), *doubled* in 65 synthetic and 34 old cultivars, and on average *46% smaller-than-doubled* in 54 wild autotetraploid species. Neither 1*x* pollen size nor 1*Cx* DNA content of diploids affected the results. Furthermore, wind-versus animal-pollination had little effect. Finally, in 16 autotetraploid-diploid pairs, the differences in pollen size were not correlated with the differences in their 1*Cx* DNA content. In sum, WGD doubles pollen volume and doubled volume persists in cultivars maintained in benign conditions, but over time in natural conditions, pollen of wild autotetraploids is strongly biased towards reversion to progenitor sizes. Since pollen downsizing after WGD was not matched by genome downsizing, pollen size reduction in young autotetraploids must largely evolve by gene regulation and expression, rather than by a direct relationship between genome size or gene dosage and cell size.

## Introduction

Polyploidy is a recurrent and rapid evolutionary process that is widespread in plants (Wood et al., 2009). The most universally observed phenotypic correlate of “pure” polyploidy is larger cell and organ size – the so-called *gigas* effects – as exemplified in young autopolyploids (Müntzing, 1936; Stebbins, 1971). *Gigas* effects are attributed to the direct biophysical impact of increased genome size (DNA mass or base-pair number) on minimum cell size and cell division rate (Bennett, 1972; Gregory, 2001; Cavalier-Smith, 2005; Francis et al., 2008; Doyle and Coate, 2019; D’Ario et al., 2021). When genome size change is by polyploidy, the epigenetic effects of larger genome size are coupled with the functional genetic consequences of doubled gene dosage and higher heterozygosity (Papp et al., 2003; Panchy et al., 2016; Husband, 2016; Birchler et al., 2021; Clo and Kolář, 2022).

In plants, most polyploid species have originated from errors of meiosis that cause whole genome duplication (WGD) in one or both gamete cell lineages (Destorme and Geleen, 2013). After fusion of its ancestral unreduced gamete(s), an autopolyploid is many cell and organismal generations removed from the original WGD(s). During stabilization and establishment of a neo-polyploid population, both the epigenetic and the genetic effects of WGD on cell size may become substantially modified by evolutionary processes. At issue are the magnitude of the pure effect of WGD on cell size, and the magnitude and direction of evolutionary change in response to WGD at different stages of polyploid establishment.

The magnitude of gigas effects varies among cell and organ types (Bannan 1948; Tsukaya, 2013; Porturas et al., 2019), ploidy level (Clo and Kolář, 2021; Becker et al., 2022), and polyploid age (Baduel et al., 2018). Furthermore, cell size is intimately linked to cell number and cell division rate, which together affect organ or tissue sizes (Šímová and Herben, 2012; Robinson et al., 2018; Simonin and Roddy, 2018; Corneillie et al., 2019; Pacey et al., 2022). Given the complexity of forces that act on cell size in multicellular tissues, the purest effects of WGD are most evident in unicellular organisms (Holm-Hansen, 1969; Glazier, 2021).

The pollen grain of flowering plants is an effectively single-celled and free-living organism. At maturity, the male gametophyte plant body comprises a single “vegetative cell” that occupies the entire volume of the mature pollen grain. Though it also forms a gametic lineage - the “generative cell” or its two sperm cell descendants - these cells are subsumed within the vegetative cell cytoplasm during early development and occupy < 0.5% of the vegetative cell volume (Russell and Strout, 2005). A mature pollen grain is an accurate indicator of the effect of ploidy on cell size because: 1) it is the same single cell or cell type in which WGD occurred, and the most common pathway to polyploidy (Ramsey and Schemske, 1998; De Storme and Geleen, 2013); 2) it is not symplastic with other cells; 3) it is terminally differentiated, arrested in G_1_ of the cell cycle (Simonini et al., 2025), and the mature size of its cytoplasm and nucleoplasm are largely controlled by gametophytic gene e*x*pression (Nelms and Walbott, 2022; Zeng et al., 2024; Li et al., 2025); 4) it has no major vacuole that distorts cytoplasm size (Murgia et al., 1987; Pacini et al., 2011); and finally, 5) its long axis is relatively stable to variation in collection conditions and is easily measured (Zuberi and Dickinson, 1985).

In the simplest scenario, WGD might result in doubled cell volume, since the cell in which non-reduction occurred is twice the size of a reduced cell, and doubled transcriptional and translational capacity might initially produce a doubled protoplasm volume. Boveri’s e*x*periments in the early 1900’s showed that unicellular sea urchin eggs doubled in volume when their chromosome numbers were doubled by polyploidy (Boveri, 1905). Boveri also generated aneuploid eggs and found that genome size (in terms of chromosome number) was correlated with cell size independently of ploidy. It is now well known that genome size (in terms of DNA mass or DNA base number) has epigenetic effects on minimum cell size, which may differ in magnitude from the effects of ploidy. Minimum cell size is largely set by genome size – by the physical space needed to accommodate extra DNA, DNA processing machinery, and associated molecules (Bennett, 1987; Cavalier-Smith, 2005; Doyle and Coate, 2019). Mechanistic hypotheses that account for the effects of both ploidy and genome size on cell size have been proposed (Amodeo et al., 2015; Schmoller et al., 2015; Amodeo and Skotheim, 2016; D’Ario et al., 2021). Though genome size limits minimum cell size, the null hypothesis for the effect of WGD on actual cell size is doubled volume (but see Adams et al., 2024).

If genome size (GS) and WGD affect cell size by the same mechanism, then the log-log correlation between pollen volume and genome size should equal 1 irrespective of the mechanism of GS change. Though some influential studies on the correlation between DNA C-value and pollen volume among species have found log-log correlation coefficients near 1 (Bennett, 1972; Lawrence, 1985 and refs therein), others have found mixed results or even no correlation (Knight et al., 2010; Moller et al., 2018; Jardine and Lomax, 2025). Comparisons of pollen size in closely related diploid and tetraploid species are similarly mixed (Stebbins 1971; Muller, 1979; Knight et al., 2010). In sum, comparative studies at the species level confound the pure effects of WGD with the effects of evolution after WGD, and so reveal little about how polyploidy actually affects the evolution of pollen size.

Size is a quantitative trait, and the genetic determinants of cell size in general are still very much an active area of research (D’Ario and Sablowski, 2019; Balachandra et al., 2022; Fung and Bergmann, 2023; Zeng et al., 2024; Li et al., 2025). In diploids, bi-directional selection e*x*periments on pollen size have found substantial heritability as well as direct impacts of larger size on seed set (Sarkissian and Harder, 2001; Lamborn et al., 2005; McCallum and Chang, 2016; Waesch et al., 2026).

Historical selection during domestication has also resulted in much larger pollen in modern diploid cultivars of both maize and rye then in their progenitors (Beadle, 1981; Waesch et al., 2025). The finding of substantial quantitative genetic variation for pollen size indicates that evolved pollen sizes in diploid species are not near a minimum, but are intermediate to their lower and upper size limits.

Pollen size in diploid species reflects an evolved compromise among its many functions, including constraints on development within the anther. Some functional aspects of pollen size are: 1) larger cells have higher metabolic vigor (Amodeo and Skotheim, 2016; Lessenger et al., 2024), and higher vigor increases pollen tube competitive ability (Kerwin and Smith-Huerta, 2000; McCallum and Chang, 2016) but shortens pollen lifespan (Williams and Reese, 2019); 2) larger cells have smaller surface-to-volume ratios which affects pollen hydrobiology and gas exchange rates (Heslop-Harrison, 1979; Ejsmond et al., 2015; Williams and Brown, 2018); 3) larger pollen size can affect cell wall thickness, increase aperture number and size, and affect wall strength (Wodehouse, 1935; Edlund et al., 2016; Lee, 1978; Whitehead and Langham, 1965); 4) larger cells have greater intracellular transport distances, cell construction costs and larger pollen tubes (Cavalier-Smith, 2005; Williams and Oliveira, 2020), potentially leading to transcription limitation (Neurohr et al., 2019; Adams et al., 2024); and 5) size-dependent pollen dispersal distances (Primack, 1978; Van Dijk and Van Delden, 1990; Aylor, 2001; Friedman and Barrett, 2009; Wilkinson, 2012; Timerman and Barrett, 2019).

Despite being unicellular with a free-living stage, pollen size is also affected by its interaction with pollen number within the developing anther and its provisioning both before dispersal and after pollination. Paternal provisioning effects are evident in negative phenotypic correlations between pollen size and number, and maternal provisioning by positive correlations between pollen size and style length (Vonhof and Harder, 1995; Hao et al., 2020; Zhu et al., 2025). In sum, the evolved sizes of pollen grains in diploid species are under strong balancing selection.

Polyploidy is a relatively instantaneous evolutionary event that could disrupt gradually evolved pollen size and its limits. After WGD, minimum cell size should increase substantially, and might fall near or even above the evolved upper size limits of its progenitor. If WGD-based size increase is e*x*cessive, it would select for reversion towards its smaller progenitor state, or perhaps only partial reversion if there are some advantages to larger size or constraints to full reversion. Taxonomic studies generally show a larger *gigas* effect on cell size in newly formed, intraspecific than in older, established autopolyploids (Kostoff et al., 1934, Tanaka, 1953; Stebbins, 1971; Muller, 1979; Butterfass, 1987; Knight et al., 2010; Tate and Simpson, 2004). Furthermore, the effect of autopolyploidy on pollen volume is stronger in transitions from haploid to diploid pollen than in transitions from diploid to higher-level pollen ploidies (Lifante, 1996; Fukuhara, 2000).

The pattern of larger effects of WGD nearer the time of WGD, is consistent with gradual weakening of epigenetic effects of GS change, or with deterministic forces that are biased to pollen size reduction over time. At the species level, the “effect of polyploidy” on pollen size is remarkably variable, which suggests that pollen size evolution after WGD is context dependent. Here I sought to understand the separate effects of WGD and subsequent earliest stages of pollen size evolution using a literature survey from mixed ploidy species.

The first questions I address are, 1) what is the magnitude of the initial effect of WGD on mature pollen size? and 2) given a large *gigas* effect, is pollen biased to size reduction after WGD in older cultivated or wild autotetraploids? Next, I considered two predictions for universal causes of pollen downsizing after WGD. First, if WGD substantially enlarges 2*x* pollen, then pollen downsizing should be stronger when pollen dispersal is directly size-dependent, as in wind-pollinated species. Secondly, if genome size directly affects cell size (Gillooly et al., 2015), and genome downsizing after WGD is common (Dodsworth et al., 2016; Baduel et al., 2022), then genome downsizing should be correlated with pollen downsizing.

I used paired comparisons of 1*x* and 2*x* pollen size in mixed ploidy species across a range of polyploid ages. First, I compared reduced 1*x* to unreduced 2*x* pollen produced in diploid anthers, and then 1*x* pollen of diploids to 2*x* pollen of their autotetraploid descendants of three ages: 1) synthetic autotetraploids (< 5 generations old), 2) cultivated autotetraploids (≥ 5 generations old), and 3) wild, long-established intraspecific autotetraploid cytotypes. I was careful to include only studies that provided data on 1*n* = 1*x* pollen size from diploid anthers and 1*n* = 2*x* pollen size from diploid or autotetraploid anthers. Butterfass (1987) used a similar approach to my first two questions, but provided no references for individual species. Here I add 61 more recent references to many published prior to 1987. I also excluded hybrids and potential allopolyploids from the data. Since the effects of WGD on size might depend on the GS and/or pollen size of the diploid progenitor, I also included 1*x* chromosome number, 1*x* genome size, and 1*x* pollen size as covariates in the analyses.

## Materials and Methods

I surveyed the literature for pollen sizes in mixed ploidy species that had both diploid and autotetraploid cytotypes. Only species in which diploids had the lowest (or near lowest, if aneuploid) chromosome number of its genus were included, such that pollen of diploids was presumed to have the 1*n* = 1*x* base chromosome number of its genus or subgenus, and pollen of autotetraploids the 1*n* = 2*x* number. Hereafter, I use 1*x* and 2*x* to refer to meiotically reduced pollen from diploid (2*x*) anthers and autotetraploid (4*x*) anthers, respectively. I also refer to meiotically unreduced pollen from diploid anthers as 2*x* pollen. For studies of diploid anthers, I avoided studies that reported cytomixis as the source of unreduced pollen (Falistocco et al., 1995), because of the high probability of aneuploidy in these. Chromosome numbers were as reported in each study, or in rare cases, taken from other literature. In this study, “autotetraploid” was considered to mean originating from diploid progenitor(s) within the same species. In the absence of genetic documentation (most studies), intraspecific tetraploids were considered to be autotetraploid if they were morphologically very similar to their diploid relative in all but cell size-related traits, and/or if meiotic or other evidence of autopolyploidy was presented. Polyploids with known interspecific contributions, either from cross-pollinations, artificially induced or spontaneous polyploidy from diploid interspecific hybrids, or natural allopolyploids, were e*x*cluded. Species with no data relevant to intraspecific polyploid parentage were also e*x*cluded.

I found 173 studies that reported pollen dimensions for both diploid and autotetraploid cytotypes within the same species. This resulted in 286 paired comparisons of 1*x* and 2*x* pollen sizes from 166 different species. Paired comparisons of 1*x* and 2*x* pollen were categorized based on the mechanism of 2*x pollen* origin and the evolutionary age of 2*x pollen* parents (Table 1).

**Table 1.** Categorization of *2x* pollen parent types.

|  | Diploid (2x) | Tetraploid (4x) |  |  |
|---|---|---|---|---|
|  | Non-reduced<sup>a</sup> | Synthetic<sup>b</sup> | Cultivar<sup>c</sup> | Wild |
| Mechanism of origin of *2x* pollen: | By errors in meiosis within diploid anther | By chemical or temperature induction | Established tetraploid cultivars | Established wild tetraploid cytotypes |
| *1x-2x* pollen size comparisons: | Same plant, often same anther | Mostly progenitor-derivative families | Mostly progenitor-derivative cultivars | Mostly closely-related, parapatric populations |
| Evolutionary age of 4x parent: |  | Generation 1-4 | Generation ≥5 | Many generations |

<sup>a</sup> 2x pollen formed *de novo* in a diploid anther, without yet having been transmitted through a sporophyte generation.
<sup>b</sup> or by spontaneous origin, if generation known.
<sup>c</sup> origin method known (spontaneous or synthetic) or unknown, but tetraploid lineage under cultivation for at least 5 generations.

Data on pollination vectors (wind, animal) was gathered from the literature, and species were classified as either animal- or wind-pollinated. Seven species known to be primarily self-fertilized or apomictic were e*x*cluded from the analysis (Supplementary data file 1). In addition, estimates of the monoploid 1*Cx* DNA content of diploids cytotypes were taken from the Plant DNA C-values Database (Leitch et al., 2019), or where possible, of both diploids and tetraploids from the primary literature. C-value estimates used either flow cytometry or the Feulgen microspectrophotometry method, which have been shown to be tightly correlated (Michaelson et al., 1991).

### Pollen size measurements

The main response variable in this study was pollen diameter, *D,* as measured by any of the methods below. Most studies reported only the diameter of the longest axis, whereas average diameter was reported in some studies with nearly spherical pollens. Some studies reported only pollen volume, and in these, *D* was calculated from:

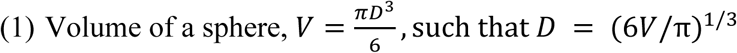

The main biological interest was the effect of ploidy on volume, and the relationship of *D* to volume might be affected by variation in pollen shape. Encouragingly, 89% of comparisons measured relatively spherical pollen in a hydrated or mostly hydrated state, whereas 32 studies reported non-spherical pollen, even after preparation. To check for effects of pollen shape, volumes were calculated from *D* or from multiple axes, if longest (*L*) and shortest (*S*) axes were measured (54 studies). For comparisons that only measured *D*, or in which *L*/*S* ≤ 1.14 (Punt et al., 2007), volume was measured as in (1) using *D* or *D* = (*L*+*S*)/2. If *L*/*S* > 1.14, then:

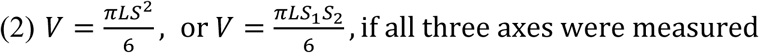

### Pollen preparation methods

Pollen preparation artifacts should be minimal, since the same method was always used to measure both 1*x* and 2*x* pollen in each ta*x*on pair and should not affect the ratio of their sizes (Müntzing, 1928). Nevertheless, since 2*x* pollen has a larger surface area, perhaps there was disproportional swelling or shrinking during fi*x*ation and/or staining. To test that hypothesis, I was able to categorize pollen preparation methods according to their potential to affect fresh pollen size in 188 ta*x*on pairs.

In the most common method, 1) fresh pollen was viewed in water or water-based viability stains such as acetocarmine, IKI, lactophenol or others, and only swollen grains that stained as viable were measured (106 ta*x*on pairs). In 2) pollen was chemically fi*x*ed before staining for viability (30 pairs). Such fi*x*atives had an aqueous component that allowed at least partial hydration of pollen before cell death (eg. < 100% Etoh, Carnoy’s, FAA). Three of these measured sizes by automatic cell sorting, after setting thresholds to remove inviable pollen and other detritus from the analysis. Finally, in 3) fresh pollen was dehydrated, either on herbarium sheets or by acetolysis (52 pairs). Dehydrated pollen was usually mounted in silicone oil or glycerin for measurement of viable pollen grains, as judged by pollen shape, not by staining – only obviously misshapen pollen grains were e*x*cluded.

Preparation methods varied strongly by pollen parent type: method 1 (fresh, hydrated pollen) was most commonly used for the three non-wild parent types (see Table 1), whereas method 3 (dehydrated pollen) was most commonly used for the wild ta*x*a. Only the wild parent type had sufficient pairs in more than one preparation category to test for a preparation effect, which was not significant (*P* = 0.208; *N* = 21, 5, 40 pairs, for methods 1-3, respectively). Therefore, pollen preparation method was not included in statistical models.

### Statistics

Pollen diameters were log-transformed and the effect of WGD on pollen size (PS) was calculated as the log-odds ratio: Δ*PS_rep_* = log_10_(2*x pollen D*) -log_10_(1*x pollen D*) for each of *N* = 288 comparisons. A species-level size effect was then estimated as: Δ*PS_species_* = arithmetic mean of Δ*PS_reps_*, which resulted in *N* = 184 species-by-pollen-parent-type comparisons. Both Δ*PS_rep_* and Δ*PS_species_* were normally distributed within each of the four pollen parent types. The same procedure was used to analyze pollen volume.

In the main analysis, I tested for difference among treatments in Δ*PS* using a mi*x*ed-model *ANCOVA*, with fi*x*ed effects of pollen parent type (0-3) and three covariates: log_10_-transformed 1*x* pollen diameter, 1*x* pollen chromosome number, and 1*Cx* value DNA content. I included species within genus as a random effect. I used the lowest *AIC_C_* as the criterion for model selection, and then Tukey’s HSD to test for differences amongst treatments. 95% confidence intervals (*CI*_95%_) were constructed to test for overlap with the doubled-volume hypothesis that mean Δ*PS* = 1.26. All statistics were done using JMP v. 15.1 (SAS institute, Cary, NC, USA).

I considered autotetraploidy within each 2*x*-4*x* intraspecific ta*x*on pair to be an independent evolutionary event. As justification for independence, Knight et al., (2010) found that evolutionary distances among species did not covary with divergences in pollen size and GS. I controlled for potential biases due to uneven ta*x*onomic sampling or higher-ta*x*on level phylogenetic effects by including a random effect of genus nested within species in each analysis.

To test whether wind- and animal-pollinated species differed in their Δ*PS* effect by parent type, I used a two-way ANOVA. Back-transformed means and 95% confidence intervals were constructed to test the hypothesis that mean Δ*PS* = 1.26. To test whether genome size differences among cytotypes were correlated with Δ*PS*, I used ordinary least squares regression of log odds ratios: Δ*GS* (log_10_ 1*Cx* genome size of autotetraploid minus log_10_ 1*Cx* diploid) on Δ*PS_species_*.

## Results

### How do autopolyploids differ from diploids in pollen size?

The final dataset included 2*x*-4*x* comparisons within 184 species, of which each of pollen parent treatments had > 30 species. In the best model, there were significant differences in the magnitude of Δ*PS* among the four pollen parent types (*P* < 0.0001), after controlling for a significant effect of chromosome number (Figure 1; Table 2). Wild species had a significantly smaller mean Δ*PS* than the two cultivated treatments (young or old), whereas diploid parents that produced unreduced 2*x* pollen displayed a significantly larger-than-1.26 effect (Figure 1). Mean Δ*PS* in young and old cultivated plants were 1.25 and 1.24, respectively (Table 2) and 95% confidence intervals overlapped the value of 1.26, the e*x*pected value if WGD doubles cell volume (Figure 1). However, the *CI_0.95_* on mean Δ*PS* = 1.158 for wild species overlapped neither 1.26 nor 1.0 (Figure 1). As a measure of precision of estimated means, the CVs calculated from standard errors were 12.2%, 5.8%, 11.1%, 6.1% for parent types 0-3, respectively. Results of full and reduced models can be found in Supplementary Tables S1A and S1B.

**Figure 1.**
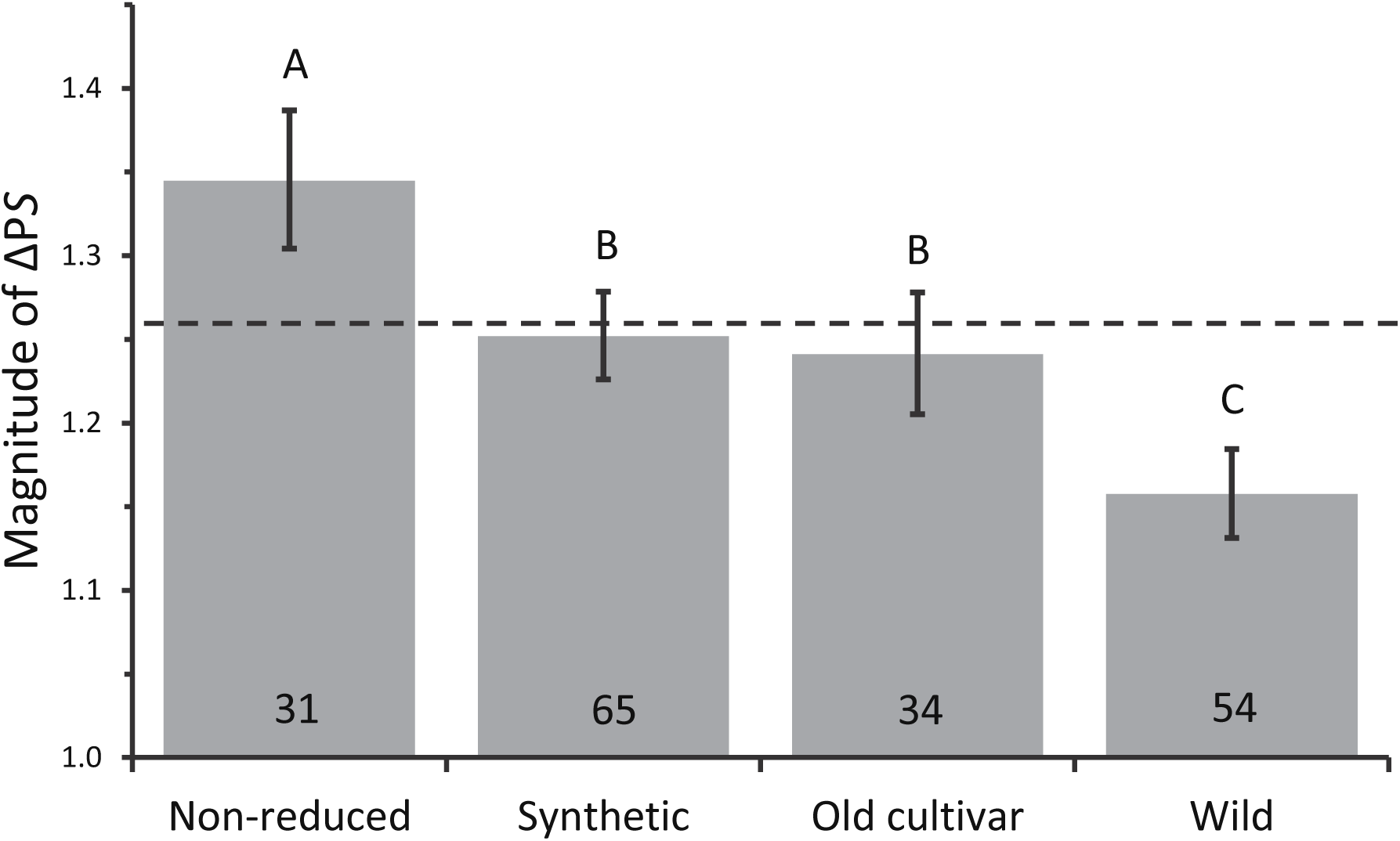
The effect of WGD on pollen size by parent type. Δ*PS* represents the ratio of 2*x* to 1*x* pollen diameter. Dashed line represents Δ*PS* = 1.26*x*, which corresponds to a 26% larger pollen diameter and doubled pollen volume in 2*x* relative to 1*x* pollen. Values are back-transformed least squares means and error bars delineate a 95% confidence interval. Bars sharing the same letter are not significantly different from each other. *N* = number 1*x*-2*x* cytotype pairs in each group, shown on graph.

**Table 2.**
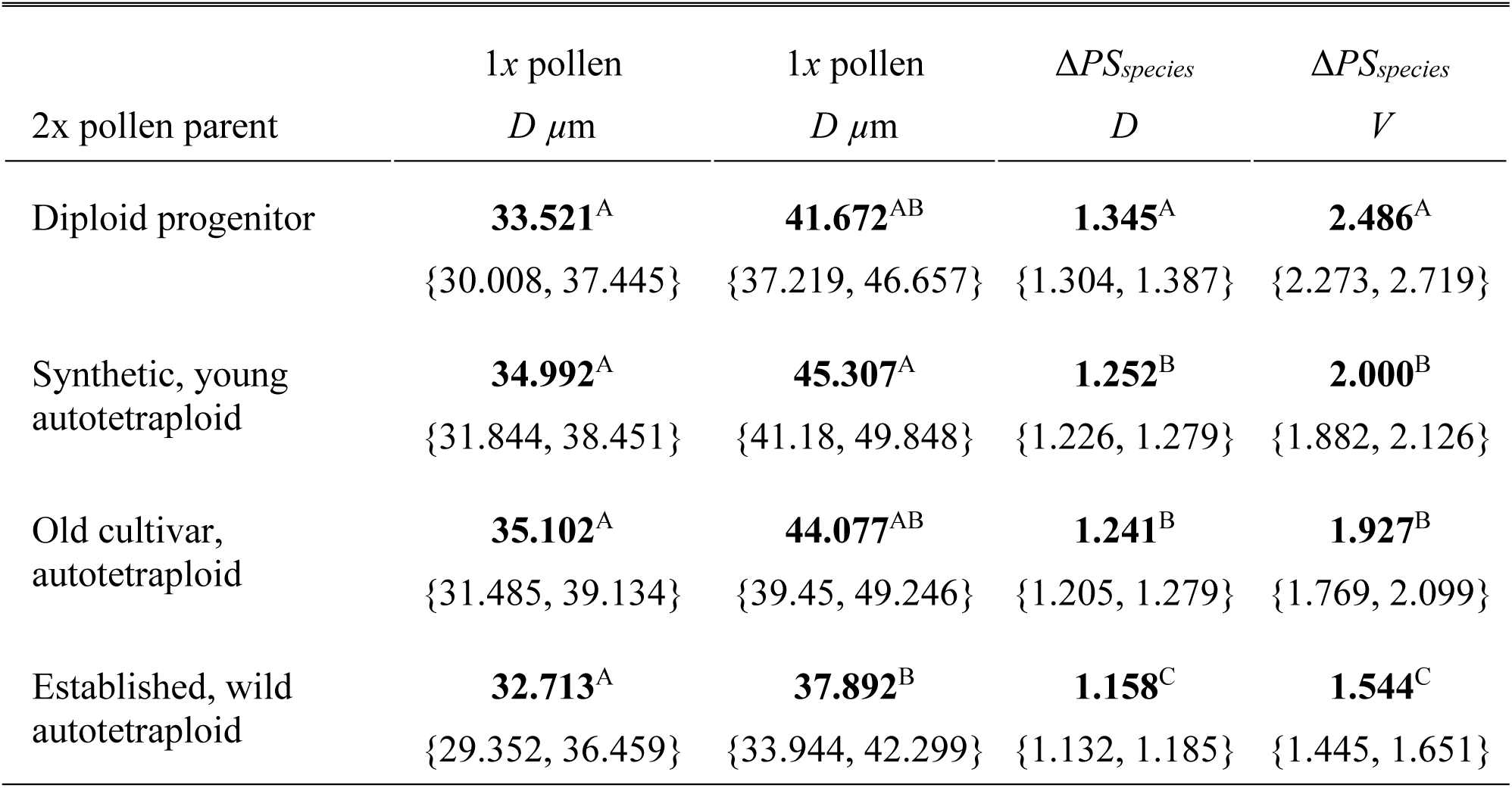
Summary statistics for 1x and 2x pollen sizes and log-odds ratios of 2x/1x pollen sizes, Δ*PS_species_*. Pollen diameters are least squares means (species nested within Genus in analysis). Δ*PS_species_* values are backtransformed means. 95% confidence intervals in brackets. Means sharing the same superscript letter within each column are not significantly different from each other. *D,* pollen diameter; *V*, pollen volume.

|  | 1*x* pollen | 1*x* pollen | Δ*PS*<sub>*species*</sub> | Δ*PS*<sub>*species*</sub> |
|---|---|---|---|---|
| 2x pollen parent | *D* *µ*m | *D* *µ*m | *D* | *V* |
| Diploid progenitor | **33.521**<sup>A</sup><br>{30.008, 37.445} | **41.672**<sup>AB</sup><br>{37.219, 46.657} | **1.345**<sup>A</sup><br>{1.304, 1.387} | **2.486**<sup>A</sup><br>{2.273, 2.719} |
| Synthetic, young autotetraploid | **34.992**<sup>A</sup><br>{31.844, 38.451} | **45.307**<sup>A</sup><br>{41.18, 49.848} | **1.252**<sup>B</sup><br>{1.226, 1.279} | **2.000**<sup>B</sup><br>{1.882, 2.126} |
| Old cultivar, autotetraploid | **35.102**<sup>A</sup><br>{31.485, 39.134} | **44.077**<sup>AB</sup><br>{39.45, 49.246} | **1.241**<sup>B</sup><br>{1.205, 1.279} | **1.927**<sup>B</sup><br>{1.769, 2.099} |
| Established, wild autotetraploid | **32.713**<sup>A</sup><br>{29.352, 36.459} | **37.892**<sup>B</sup><br>{33.944, 42.299} | **1.158**<sup>C</sup><br>{1.132, 1.185} | **1.544**<sup>C</sup><br>{1.445, 1.651} |

The mean (median) base 1*x* chromosome number for 167 species was 10.7 (10), ranging from 5 to 24 with one outlier (*Adansonia,* 1*x* = 44; Baum and Oginuma, 1994; Udeh, 2017), and one uncertain count in *Begonia* (see Supplementary data file 1). The log-log slope of 1*x* chromosome number on Δ*PS* was 0.046, which indicates that as chromosome number increased, there was a small, but proportional increase in Δ*PS*. Δ*PS* increased by 7.5% over the range of 1*x* = 5 to 24 chromosomes in this study. There was information for 1*Cx* genome size in 133 species (see Supplementary data file 1). 1*Cx* DNA content and its interaction with 1*x* chromosome number were dropped from the full model (*P* = 0.986, 0.329, respectively; Supplementary Table S1B).

Since pollen parent treatments comprised different sets of species, it is possible that evolved pollen sizes of diploids in the wild were smaller or larger than in other parent types, and that the difference might affect Δ*PS.* Importantly, there was no evidence that Δ*PS* scales to the size of 1*x* pollen (covariate *P* = 0.553) or that it scales differently in different populations (interaction effect, *P* = 0.363; Supplementary Table S1B). Furthermore, 1*x* pollen diameters did not differ between treatments (*P* = 0.64). Mean pollen diameters were: 33.5, 35.0, 35.1, and 32.7 *µ*m for parent types 0-3, respectively (Table 2). On the other hand, 2*x* pollen diameters of wild species were smaller than those of synthetic autotetraploids (Table 2). Thus, the smaller Δ*PS* value in wild species was due to reduced 2*x* pollen diameter, not to increased 1*x* pollen diameter.

The results for pollen volumes were substantially similar to those for pollen diameters, even though pollen shapes varied and volume was calculated by different methods (Table 2). There were significant differences among 2*x* pollen parents (*R^2^_adj_* = 0.31; *P* < 0.0001), after accounting for a positive effect of chromosome number (*P* = 0.017). The confidence intervals of newly synthesized and old cultivar autopolyploids overlapped 2.00, or doubled volume, whereas wild species had significantly smaller-than-doubled pollen volume.

To check whether volume measurement method affected results, volumes measured by formula 1 (spheroidal pollen) versus formula 2 (non-spheroidal pollen) were averaged separately within each species and 2*x* pollen parent type. A two-way ANOVA indicated differences among 2*x* pollen parent types (*R^2^_adj._* = 0.27; *P* < 0.0001), but no differences in volume measurement method (*P* = 0.886) and no evidence for a parent x method interaction (*P* = 0.192). Sample sizes for non-spheroidal pollen were *N =* 4, 10, 2, and 8, respectively.

### Do wind pollinated species have smaller ΔPS than animal pollinated species?

I was able to categorize 177 species for pollination vector, of which 37 were wind- and 140 animal-pollinated (the other 7 were autogamous). A two-way ANOVA found a significant parent type by pollination mode interaction (*P* = 0.007; Supplementary Tables S2A, S2B). In separate one-way ANOVAs on each pollen parent type (Figure 2), pollination modes did not differ in Δ*PS* for diploid and newly synthesized autotetraploid parents (*P* = 0.281, 0.981, respectively). In older cultivars and established wild ta*x*a, Δ*PS* of wind pollinated autotetraploids was marginally smaller than in animal pollinated species (*P* = 0.003, 0.083, respectively). In wild species, the 95% confidence intervals of both wind- and animal-pollinated species do not overlap 1.26, whereas in cultivars, Δ*PS* of animal-pollinated species is consistent with 1.26. Δ*PS* was smaller than 1.26 in both wind-pollinated cultivars and wild species.

**Figure 2.**
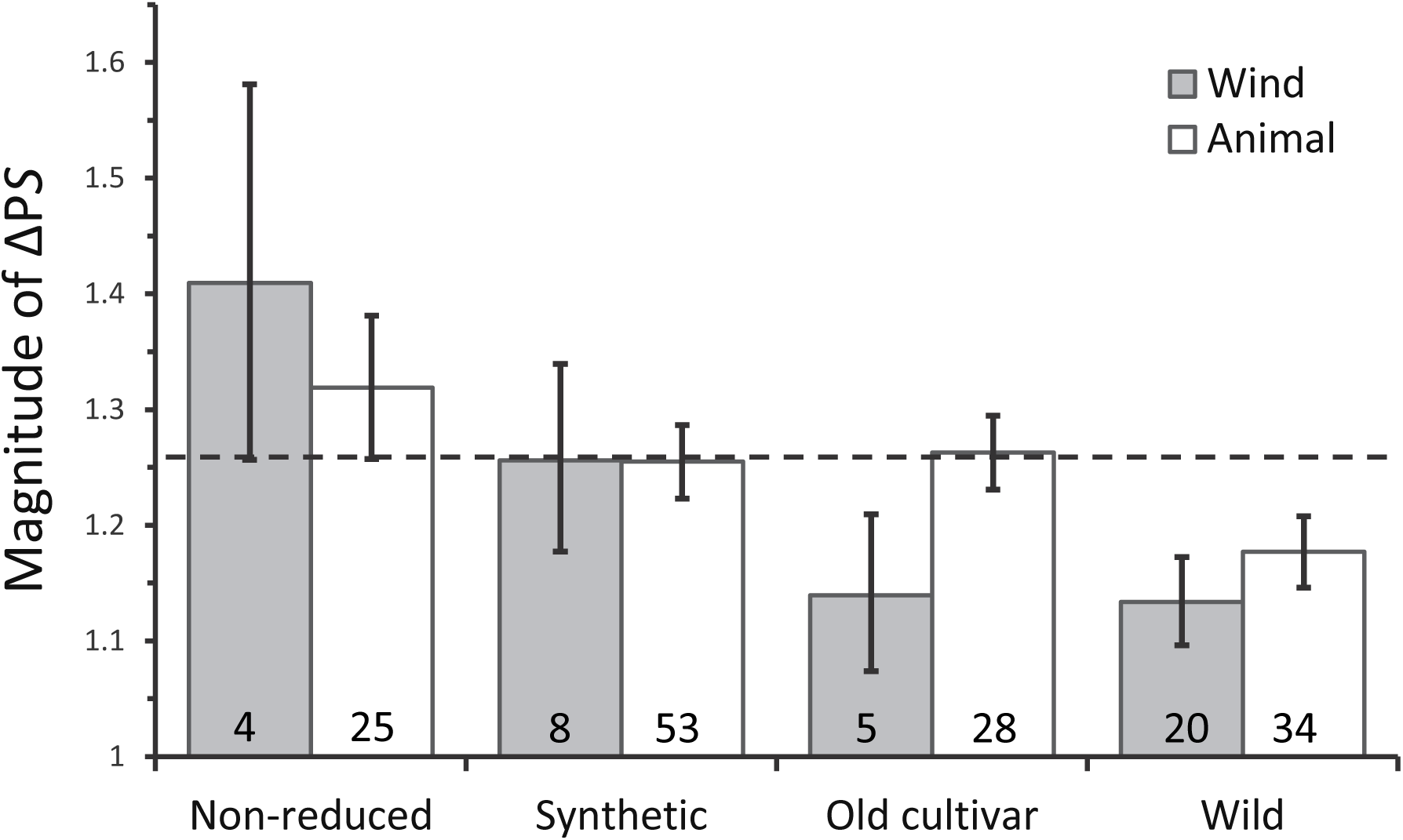
The effect of WGD on pollen size by pollination type. Δ*PS* represents the ratio of 2*x* to 1*x* pollen diameter. Dashed line represents Δ*PS* = 1.26*x*, which corresponds to a 26% larger pollen diameter and doubled pollen volume in 2*x* relative to 1*x* pollen. Values are back-transformed least squares means with error bars delineating a 95% confidence interval. *N* = number 1*x*-2*x* pollen cytotype pairs in each group, shown on graph.

### Does genome downsizing explain the smaller ΔPS of wild species?

The 1*Cx* (monoploid) genome sizes of 2*x* and 4*x* cytotypes should be the same at the time of WGD, but subsequently, genome downsizing might occur. DNA 1*Cx*-value data was available for both cytotypes in 16 wild species (refs in Leitch et al., 2019 and Supplementary data file 1). In this group, 4*x* cytotypes had a slightly lower mean 1*Cx* value than 2*x* cytotypes (*P* = 0.061; one-tailed matched pair test). Pollen downsizing also occurred in the autotetraploids. The back-transformed LS mean Δ*PS* = 1.143, CI_95%_ {1.110, 1.178}, which is very near that of the full set of wild species (Table 2).

If pollen size is mechanistically linked to genome size, then in wild species the magnitude of post-WGD genome downsizing should scale with the magnitude of 2*x* pollen downsizing. There was no evidence for a positive correlation between Δ*GS* (autotetraploid log_10_ 1*Cx* minus diploid log_10_ 1*Cx*) and Δ*PS* (Pearson correlation, *r* = -0.384; *P* = 0.142; *N* = 16; Figure 3). Fourteen of 16 autotetraploids had >20% smaller 2*x* pollen size than expected at WGD, whereas only three had >20% smaller genome size than expected at WGD (Figure 3).

**Figure 3.**
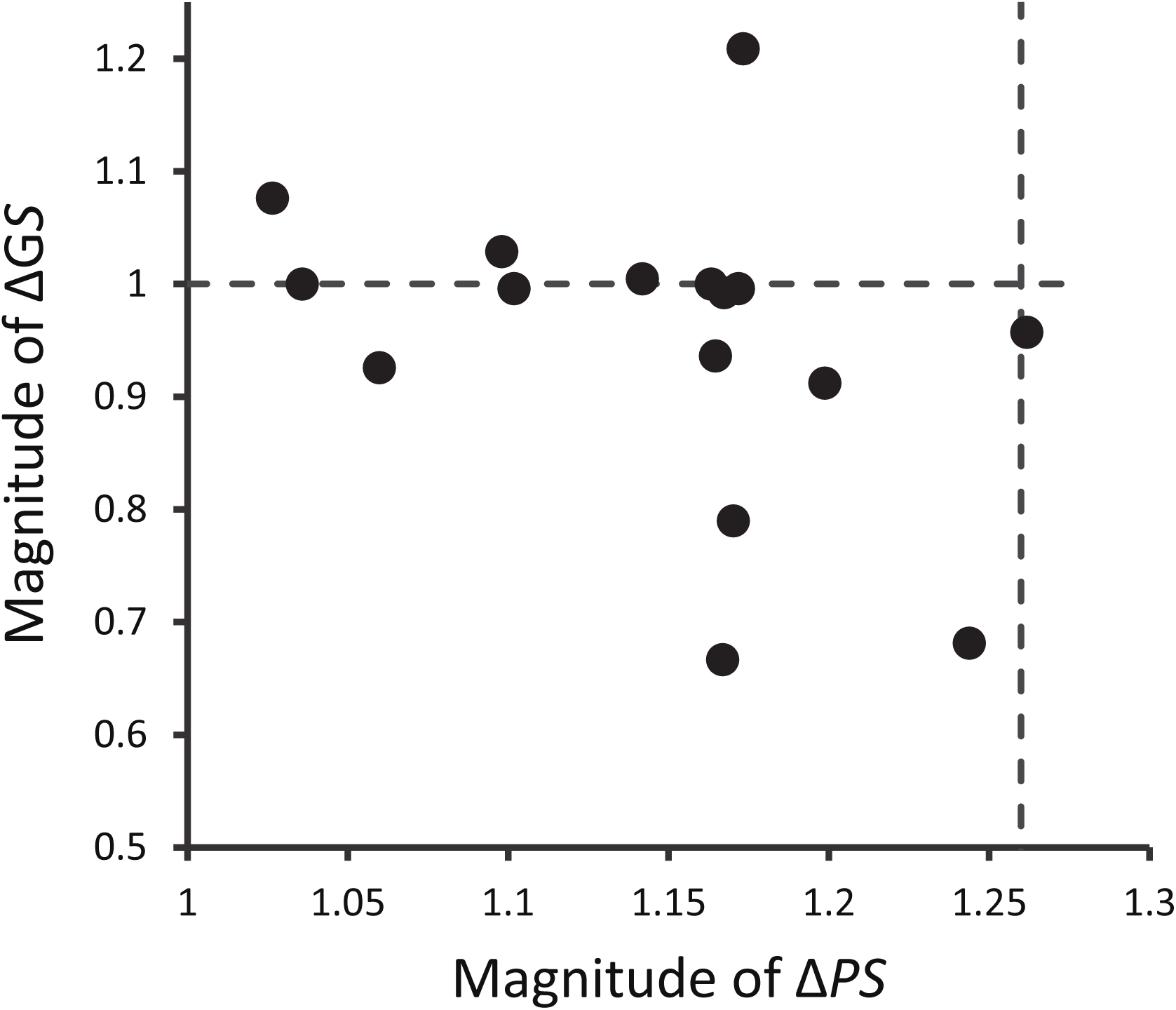
The relationship between the effects of WGD on 1*Cx* genome size and on pollen size in wild species. Δ*PS* represents the ratio of 2*x* to 1*x* pollen diameter and Δ*GS* the ratio of 2*x* to 1*x* monoploid (1*Cx*) genome size. At inception of WGD, Δ*PS* and Δ*GS* are expected to be 1.26 and 1.0, respectively (dashed lines), whereas if full reversion to progenitor sizes occurs, then Δ*PS* = 1.0 and Δ*GS* = 0.5. Values have been back-transformed from log-odds ratios used in the analysis.

## Discussion

Genome size change by polyploidy has large effects on cell size, but both the magnitude and evolutionary duration of *gigas* effects have been debated. Where minimum cell size limits are set by genome size, upper limits may be set by development, physiology, and the environment.

Instantaneous change in pollen size associated with WGD might cause 2*x* pollen to e*x*ceed optimal or upper cell size limits of its evolved 1*x* progenitor. On the other hand, reversion to progenitor size might not be possible if the new minimum size limit of 2*x* pollen after WGD is above the previously evolved optimal size of 1*x* pollen. The results here show that 2*x* pollen volume in the first few autopolyploid generations after WGD is on average double that of 1*x* pollen of its intraspecific progenitor. Secondly, evolved 2*x* pollen of wild neo-autotetraploids is much smaller than doubled 1*x* pollen volume, but still larger than 1*x* pollen. These results indicate that after WGD in the wild, 2*x* pollen size is biased to rapid reduction. Dispersal size may drive reduction, but pollen downsizing was similar in magnitude in both wind- and animal-pollinated wild autotetraploids, which suggests other forces are also causing reduction. Genome size reduction could cause pollen downsizing, but most wild 4*x* cytotypes had substantial pollen downsizing without genome downsizing. Thus, early phenotypic evolution in autopolyploids must occur more by recombination of their larger genomes coupled with changes in gene expression and regulation than by point mutations and deletions.

Variability in the degree of pollen downsizing, may also be due to variable minimum cell size limits and the nature of size-number trade-offs during establishment.

### The “pure” effect of WGD in diploid anthers is at least doubled pollen volume-

When 2*x* pollen is formed via meiotic non-reduction within a diploid anther, the volume of the 2*x* microspore after cytokinesis is roughly the volume of two of the four 1*x* microspores, or about half the original volume of the microsporocyte (Lee et al., 2026). At inception, a 2*x* microspore is double in size of a 1*x* microspore (White and Davidson, 1979). If growth and development of both are directly proportional, then a mature 2*x* vegetative cell should be double the volume of its 1*x* progenitor. Yet, 2*x* pollen in diploid anthers was on average nearly 2.5-fold larger than 1*x* pollen in the same anther. Though variance among studies of 2*x* pollen in diploid anthers was high, 2*x* pollen was more than double the volume of 1*x* pollen in 65% of comparisons.

There are several reasons why 2*x* pollen formed by non-reduction might appear to be more than double the size of 1*x* pollen. First, unreduced 2*x* pollen is almost always identified by its large size alone, and researchers generally adopt a threshold diameter to positively identify pollen as 2*x*. Since the diameter of a sphere of doubled volume is 126% of the original, Darlington (1937) recommended a 125% threshold. More recent studies have generally recommended a more conservative value of 130 to 140% (Zlesak et al., 2009; Gao et al., 2019), which is consistent with the mean 134% effect of WGD found here. At the same time, non-reduction at both meiosis I and II can sometimes produce 4*x* pollen, which is even larger than 2*x* pollen. Cryptic cytomixis events can also result in right-skewed distributions of 2*x* pollen size due to the inclusion of both aneuploid and 4*x* pollen (Falistocco et al., 1995). Thus, 2*x* pollen size can be slightly overestimated by thresholding out some smaller 2*x* pollen and by inclusion of larger pollen with > 2*x* chromosome numbers. As an e*x*ample of the thresholding effect, in diploid *Lolium perenne*, Sala et al., (1989) used a 126% threshold to identify “big” (non-reduced) pollen visually and reported it to be much larger than double the volume of 1*x* pollen. But a subsequent study on the same species used statistical methods to separate overlapping size distributions of pollen within anthers, and the resulting means were very near to the 1:2:4 volume ratio expected for 1*x*, 2*x* and 4*x* pollen volumes (Jansen and Den Nijs, 1993). Sometimes distributions have little overlap and thresholds are not used. Beadle (1981) treated diploid teosinte plants with colchicine and found that abnormally large, presumably 2*x*, pollen grains were on average exactly 26% larger in diameter than 1*x* pollen from the same anther, or from untreated plants

Other methods have been used to score “big” pollen as non-reduced, such as the unique shapes, increased aperture numbers, and/or enlarged nuclei of 2*x* pollen (Dewitte et al., 2009). Flow cytometry can also be used to verify that “enlarged nuclei” of large, putative 2*x* pollen grains actually have doubled DNA C-value (Destorme et al., 2013). For example, Destorme and Geleen (2011) found that non-reduced 2*x* pollen from diploid *Arabidopsis Col-0* were 24% larger in diameter than 1*x* pollen, which is close to the 28% larger pollen in known autotetraploids relative to diploids.

Some studies collected data on the frequency of non-reduction in meiocytes and were able to count the number of non-reduced and reduced nuclei to estimate the expected percentage of mature reduced 1*x* and non-reduced 2*x* pollen. In *Ziziphus jujuba,* Xue et al., (2011) found the frequency of non-reduction in meiocytes (5.7%) was not significantly different than that of putative 2*x* pollen (6.2%). That 2*x* pollen was recognized using a high threshold of 150% suggests that 2*x* pollen is sometimes truly larger-than-double in size. In fact, *2x* pollen of an old 4*x* cultivar of *Z. jujuba* was 146% larger than its diploid progenitor cultivar (Wang et al., 2019).

There may be biological reasons for non-reduced 2*x* pollen to be > 2-fold larger in volume than 1*x* pollen in diploid anthers. One biophysical explanation is in meiocyte size – in species with simultaneous cytokinesis (most eudicots), smaller microsporocytes are more likely to undergo non-reduction due to a higher chance of spindle interaction during meiosis II (Yi et al., 2023). If the resulting non-reduced microspore(s) acquires half the original protoplasm and relative sizes are maintained during growth and development, then mature 2*x* pollen should be smaller, not larger, than double the size of 1*x* pollen.

But there are reasons that size differences might arise during cell growth. During microsporogenesis, the callose wall seals off the microsporocyte and meiotic products from sporophytic influence until its dissolution. 2*x* microspores are formed by failure of cytokinesis between two meiotic products followed by the envelopment of the two chromosome sets in a single nuclear envelope. These energetic activities are governed by sporophytic resources garnered before the callose wall was formed (Marchant and Walbott, 2022), and if equal partitioning occurs, then a neo-2*x* microspore should have twice the resources as two 1*x* neo-microspores. But the neo-2*x* microspore has also saved energetic resources by not having built the equational plasma membrane and wall, and using less-than-double the amount of nuclear membrane, plasma membrane and intine material to grow into a mature pollen grain. A spherical 2*x* cell or nucleus has appro*x*imately 20% less surface area than that of two 1*x* spheres of half volume. Theoretically, the higher metabolic activity of larger initial cell size coupled with its lower costs of growth might result in building a cell of larger-than-double volume relative to, or even at the expense of, its two 1*x* monad brothers.

Polyploidy has been invoked as a mechanism for selection on cell size (Gregory, 2001; Cavallier-Smith, 1978). Given the advantages of size and gene dosage in unreduced 2*x* pollen, the post-pollination process of intra-individual pollen competition (Mulcahy and Mulcahy, 1987) might originate in the anther. Dewitte et al., (2009) found that despite low numbers of unreduced 2*x* pollen compared to 1*x* pollen within anthers, 2*x* pollen must often outcompete 1*x* pollen for fertilizations - in diploid-diploid crosses from such males, more seeds were triploid than expected based on the frequency of non-reduced pollen.

Competition could occur within the anther by differential provisioning of 2*x* microspores, or through greater resource acquisition of 2*x* microspores. Though the initial stages of microspore growth are governed largely by sporophytic transcripts and proteins (Marchant and Walbott, 2022), gametophytic gene e*x*pression predominates at the first mitosis that forms the vegetative and generative cells (Nelms and Walbott, 2022). This means that gametophytic gene expression governs the major growth phase of the vegetative cell (Hafidh et al., 2016) when diploid, unreduced pollen is in direct competition with haploid, reduced pollen for resources. Whereas theory indicates equal provisioning of spores confers higher fitness to the spore parent (Haig, 2010), formation of 2*x* pollen may be a mechanism for avoiding equal provisioning. When 2*x* pollen is formed by non-reduction in a diploid anther, its genetic or epigenetic size has yet to be transmitted by fertilization through a sporophyte generation, where maternal contributions to 2*x* size might come into play, and where all pollen in the anther is 2*x*. Interestingly, when autotetraploidy is artificially induced on the meristems of diploid plants (the C_0_ generation), such that there are both diploid and tetraploid flowers on the same “mixoploid” plant, the median 2*x* pollen volume from autotetraploid anthers was 211% that of 1*x* pollen from diploid anthers (Butterfass, 1987).

### The “pure” effect of WGD is transmitted as doubled pollen volume in young tetraploid sporophytes

2*x* pollen produced by autotetraploid anthers in first or early generation autotetraploids was on average almost e*x*actly double the volume of pollen from diploid progenitors. Whereas the synthetic or spontaneous autotetraploids have nearly the same genotypes as their diploid parents, older animal-pollinated cultivars also maintained exactly doubled volume of 2*x* pollen. Of the four pollen parent types, the most precise estimate of mean Δ*PS* was for young synthetic autotetraploid and diploid progenitor pairs, which had a *SE*-based *CV* of 5.8%. Variation in the effect size of WGD, the log-odds ratio Δ*PS,* was also normally distributed, consistent with stochastic variation around a mean.

In artificial or spontaneous autotetraploids, microsporocytes (pollen mother cells, or meiocytes) in early or late meiosis were close to double the volume of diploid progenitor microsporocytes (Belling and Blakeslee, 1923; Sax and Sax, 1937; Bannan, 1948). In *Tradescantia* autotetraploids, microsporocytes in meiosis were 19% longer, and mature pollen 17% longer, than in diploids (Sax and Sax, 1937). In 2*x* and 4*x Taraxacum*, 1*x* and 2*x* microspores increased in diameter by 5.56-fold and 5.34-fold, respectively, during pollen maturation (Bannan, 1948). The closely comparable growth patterns, from microsporocyte to microspore to mature pollen, suggests that mature pollen size is transmitted largely by the initial properties of the microsporocyte and its partitioning into microspores, not by differential growth.

Microsporocyte sizes within an anther are relatively homogeneous (Bannan, 1948; Yi et al., 2023; Lee et al., 2026), as they derive from a symplastic sporogenous tissue and are synchronized in developmental stage (Heslop-Harrison, 1966; Whelan, 1974; Magnard et al., 2001; Blackmore et al., 2007; Li et al., 2015; Marchant and Walbott, 2022). Since sporogenous tissue, which gives rise to microsporocytes, is an early determinant of anther size, and mature pollen size is limited by anther locule size, both microsporocyte and pollen sizes are tightly regulated (Yi et al., 2023). Variation on the small side can cause errors in meiosis I because space constraints can allow spindles to interact, leading to non-reduction (Yi et al., 2023, but see Lee et al., 2026). Variation on the large side can lead to reduced pollen numbers, affecting male function. Viable pollen within an anther is also very uniform in size, and CVs are usually in the range of 1-5% (Vonhof and Harder, 1995), and not affected by the usual sources of variation in cell size, such as vacuole size or constraints of development within multicellular tissues. Thus, the purest, heritable effect of WGD on pollen size should be most apparent in early generation autotetraploids.

Studies on inbred lines have reported variation among genotypes in the magnitude of Δ*PS*, but the averages were still close to 1.26: 1.25 in *Zea mays* (Southworth and Pfahler, 1992), and 1.276 in *Arabidopsis* (Altmann et al., 1994). Tsukaya (2013) also found Δ*PS* from three mutant lines varied on both sides of a wild type value of 1.279, and concluded, “Ploidy level may affect cell volume, but the magnitude of cell expansion is likely under control of genetic pathways.” Given the consistency of the results here, which center on an effect size of doubled volume in the first few generations after WGD (from 106 estimates in 65 species), I conclude that in taxa that have evolved their haploid pollen size naturally (i.e., “wild type”), the initial effect of WGD is to double pollen volume.

### After WGD, widespread convergent evolution to smaller 2x pollen size

Intraspecific 2*x* pollen from wild autotetraploids was significantly smaller than doubled volume of 1*x* pollen of diploid progenitors. Butterfass (1987) found that pollen volume in 44 wild species had Δ*PS* = 1.67, which is near the value of 1.54, from 54 species, which included more recent studies. I also found that the smaller apparent 2*x* pollen size increase in wild than in synthetic autotetraploids was not due to different starting points – diploid progenitors of synthetic and wild autotetraploids had similar mean 1*x* pollen sizes, but different 2*x* pollen sizes. The best explanation for these facts is that wild autotetraploids have convergently evolved smaller pollen sizes after WGD.

Given the consistency of doubled volume in synthetic autotetraploids, I infer that on average wild intraspecific autotetraploids have lost some 46% of the pollen volume gained from WGD. But 46% is an underestimate, since the majority of studies of diploid-autotetraploid pollen sizes were based on the taxonomic value of large size differences in the first place. In studies of wild populations, minimal differences are rarely reported, and when reported, are rarely measured (eg., Baldwin, 1936; Soltis, 1984). There were four cases of 2*x* pollen volume that had reverted to within 10% of their 1*x* progenitor size among 54 independent wild autotetraploid origins (and another below).

Only two studies included diploid, synthetic neo-4*x*, and wild 4*x* plants (a third was published after analyses were done; Westermann et al., 2024). In these, the magnitudes of both the gain by WGD and the loss after WGD can be estimated in the same species. 2*x* pollen in *Chamerion angustifolium* underwent a 77% gain in volume by WGD, followed by 15% loss (Baldwin and Husband, 2001); in *Arabidopsis arenosa,* a 98% gain, followed by an 89% loss (Westermann et al., 2024); and in *Plantago media,* a 111% gain followed by an 87% loss (Van Dijk and Van Delden, 1990).

Importantly, in *A. arenosa*, there were two independently originated autotetraploid populations, and these had parallel volume losses of 77% and 101% (Westermann et al., 2024). These studies support the conclusion of substantial biased evolution to smaller pollen size after WGD.

Bias to reduction suggests that instantaneous doubling of pollen volume disrupts size-dependent functions and favors a reversion to progenitor size. One mechanism for reversion is through genome downsizing, which would affect both sporophytic and gametophytic cells. For example, stomatal guard cells exhibit a similar pattern as pollen – near doubled volume in synthetic autotetraploids and less-than-doubled volume in wild autotetraploids (Butterfass, 1987). Genome downsizing is hypothesized to be directly involved in reducing leaf cell sizes for more efficient performance (Hodgson et al., 2010; Simonin and Roddy, 2019). If genome size is directly involved in setting minimum cell size, then shifts to smaller genome sizes should relax biophysical constraints, allowing pollen size reduction as well. Genome downsizing should scale with pollen downsizing in each wild intraspecific autotetraploid, resulting in a positive linear log-log correlation (Fig. 3). But there was no correlation, and instead, a pattern of large reductions in 2*x* PS but few reductions in autotetraploid GS. It appears that pollen size evolves more rapidly than GS after WGD, consistent with studies that show the rate of genome downsizing in intraspecific autopolyploids is very slow (Greilhuber and Leitch, 2013; Wang et al., 2021). Though WGD doubles both pollen volume and GS, after WGD genome size and pollen size evolution are clearly decoupled.

One explanation for why pollen downsizing should be so rapid is that reversion of epigenetic effects of WGD is easier to accomplish and can occur faster relative to the timeframe of change by genetic mutation, pseudogenization, and genome downsizing. The relative stability of genome size in young tetraploids suggests that rapid changes in gene expression and regulation after WGD (eg., Srikant et al., 2024) are the earliest mechanisms for pollen downsizing, a case of phenotypic evolution preceding genic evolution (West-Eberhard, 2003). Selection on pre-existing variation and recombination could also contribute to rapid downsizing (eg., Hollister et al., 2012). Pollen size in most species has substantial additive or broad sense genetic variation, and experimental selection for smaller pollen size in diploids was effective (Sarkissian and Harder, 2001; Lamborn et al., 2005; McCallum and Chang, 2016). Autopolyploidy elevates allelic diversity and mutation rates in individual sporophytes (Parisod et al., 2010), and especially in diploid relative to haploid gametophytes (Williams, 2021). For these reasons, gametophytic selection on recombinants and new, but rare mutations might also contribute to size reduction in 2*x* pollen.

The striking fact of independent, repeated shifts to smaller size in most intraspecific autotetraploids suggests that pollen enlargement by WGD was generally maladaptive. Pollen size has a more direct impact on dispersal distance when pollen is dispersed by wind than by animals (Welsford et al., 2016; Zhu et al. 2026). As a result, wind-pollinated neo-tetraploids should experience stronger selection for smaller 2*x* pollen size than animal-pollinated neo-tetraploids (Wodehouse, 1935; Van Dijk and Van Delden 1990; Bretagnolle and Thompson 1995; Knight et al., 2010). 2*x* pollen in wind-pollinated old cultivars was smaller-than-doubled, whereas animal-pollinated old cultivars had doubled 2*x* volumes, similar to synthetic neo-autotetraploids. It may be that cultivation in benign environments or younger evolutionary age allowed animal-pollinated autotetraploids to retain their doubled size (compared to wild autotetraploids that did not). Wind-pollinated old cultivars had smaller-than-doubled 2*x* pollen size, but to the same degree as wind-pollinated wild autotetraploids. Selection on dispersal distance may have reduced size in both, but in the wild, 2*x* pollen of animal-pollinated autotetraploids experienced a similarly large loss. If wind-pollination plays a role in 2*x* pollen downsizing, some other cause must also affect size independent of pollination vector.

One obvious possibility in wild environments is stronger selection for smaller size due to greater resource or pollen limitation. Pollen metabolic demands are among the highest of any plant cell type (Rounds et al., 2011), and wild neo-polyploids might be unable to completely provision pollen of larger size. Nutrient limitation might have affected some neo-polyploids, but polyploid plants are also increasingly being found to compensate for their larger sizes via greater resource acquisition ability (Bomblies, 2020; Šmarda et al., 2023; Sidhu et al., 2025). When resource limitation does occur, it is more likely to affect the whole plant, resulting in fewer or smaller flowers, so the question is whether plant downsizing results in fewer pollen numbers or smaller pollen sizes.

In early tetraploid establishment, ovules are predominantly fertilized by haploid, not diploid, pollen (Levin, 1975), and since most autotetraploids are outcrossing (Husband et al., 2008), tetraploid-tetraploid matings only occur in rare dispersals of 2*x* pollen to 2*x* ovules. Thus, the size and number of pollen grains play an outsized role in reproductive success during early polyploid establishment. The metabolic benefits of larger 2*x* pollen size and doubled gene dosage are consistent with 2*x* male gametophyte physiology: faster meiosis and pollen development of autotetraploids than diploids (Bennett and Smith, 1972; Williams and Birmley, *manuscript available*); the faster than expected pollen tube growth rates (PTGRs) of intraspecific autotetraploids and polyploid species (Reese and Williams, 2019; Williams and Oliveira 2020; Westermann et al., 2024); and the angiosperm-wide evolution of faster PTGRs in polyploid compared to diploid species (Reese and Williams, 2019). But the effects of WGD on size, vigor and performance might be less important initially than the effects of WGD on pollen number.

Pure polyploidy increases cell and organ sizes, but cell division rates in tissues are either unchanged or reduced in compensation (Greilhuber and Leitch, 2013; Robinson et al., 2018). Pollen develops in an anther, and pollen precursor cells in the core tissue of a developing anther play a role in determining anther size. Thus, autopolyploid pollen production is expected to be similar or lower than production in diploid progenitors. There is intense selection on pollen viability during the first few generations of autopolyploid formation (Stebbins, 1971; Ramsey and Schemske, 2002; Bomblies, 2023). Fertility restoration involves both the stabilization of meiosis, as well as adjustments to pollen phenotypes (Westermann et al., 2024), which reverts viability towards diploid progenitor levels. Selection for higher fertility might also affect pollen production. For example, tetraploid species have evolved greater pollen production per anther than their nearest congeneric diploid relatives in 20 of 44 comparisons (Williams et al., 2026). Importantly, 2*x* pollen size reduction after WGD may be the enabling factor in increased pollen production per anther - at the intraspecific level, pollen size reduction after WGD was correlated with increased pollen production in 3 of 5 wild autotetraploids (Oliveira et al., 2022; Baldwin and Husband, 2011; Williams et al., 2026). Pollen number per flower increases with pollination uncertainty (Harder and Johnson, 2023), and therefore is likely to be important in mate limited situations (Cunha et al., 2022), such as in newly establishing populations of outcrossed neo-autopolyploids.

In sum, post-WGD evolution of pollen size is context-dependent and may depend strongly on how pollen size-number trade-offs are resolved at different stages of early polyploid evolution. Fitness gains through increased 2*x* pollen production (coupled with decreased 2*x* pollen size) must be important during early autotetraploid stabilization and establishment, whereas retaining some of the WGD volume-doubling along with doubled transcription capacity might reinforce 2*x* pollen competitive ability on both diploid and autotetraploid styles. Male function must play a large role in early establishment of outcrossing polyploids, but models rarely, if ever, consider the joint effects of pollen size and pollen number on fitness outcomes (see Osterman et al., 2025). Excessive pollen or cell size may be maladaptive, but retention of some size increase along with WGD is clearly a common pattern in early autopolyploid evolution. After all, almost all wild autotetraploids reviewed here retained larger-than-progenitor 2*x* pollen sizes, and more than a quarter still had pollen within 10% of doubled volume or larger.

## Author Contributions

The author designed the study, conducted all research, performed all analyses, and wrote the manuscript.

## Acknowledgments

The author thanks B. O’Meara and H. Yomai for useful discussions.

